# Dynamic transcription pre-initiation complex assembly governs initiation efficiency

**DOI:** 10.1101/2025.05.07.652662

**Authors:** Nayem Haque, Robert A. Coleman

## Abstract

Transcription initiation is a highly regulated process that determines gene expression outcomes^1,2^, yet the dynamics of initiation and the mechanisms governing efficiency remain poorly understood. Here, we combine endogenous tagging of human RNA polymerase II (Pol II) and TFIID with simultaneous live-cell, multi-color single-molecule imaging to quantitatively map Pol II behavior during transcription initiation and early elongation. Using GRID (Genuine Rate Identification) analysis, we resolved four distinct kinetic populations of chromatin-bound Pol II. The dynamics of Pol II populations reveal that initiation is highly inefficient, with over 94% of Pol II molecules dissociating within tens of seconds. Kinetic partitioning of Pol II dwell times enables quantification of proximal pausing, which is globally sensitive to CDK9 inhibition. Single-cell analysis uncovers substantial heterogeneity in initiation efficiency and pausing across individual cells. Colocalization of Pol II with TFIID is associated with higher initiation efficiency and reduced promoter-proximal pausing compared to global Pol II. Further dissection of Pol II–TFIID assembly pathways reveals that canonical assembly, where TFIID binds first, is linked to inefficient initiation and frequent pausing. In contrast, non-canonical assembly, where Pol II binds first followed by TFIID, supports more efficient initiation with lower pausing. Together, these findings establish that transcription initiation efficiency is shaped by both the kinetic stability of Pol II engagement and the temporal order of pre-initiation complex assembly, providing a new framework for understanding dynamic gene regulation in vivo.

## Introduction

Transcription initiation is a highly regulated and dynamic process that governs gene expression across all living cells^1,2^. It begins with the binding of TFIID to the core promoter, followed by recruitment of RNA polymerase II (Pol II) and assembly of the pre-initiation complex^3-5^. The transition of Pol II from promoter binding to productive elongation represents a major regulatory checkpoint and rate-limiting step in transcriptional activation, influencing cell identity, developmental programs, and responses to environmental stimuli^6^. Although the core molecular machinery of initiation has been well characterized through biochemical and structural studies, how initiation efficiency is regulated within the native nuclear environment remains poorly understood.

Previous FRAP-based studies using locus-specific reporters and ensemble measurements have suggested that transcription initiation is highly inefficient, with most Pol II molecules dissociating during early stages of promoter engagement^7,8^. However, these approaches rely on engineered systems and population averages, which may not reflect general features of endogenous transcription. Moreover, the extent to which transcription factor dynamics and pre-initiation complex assembly pathways influence Pol II initiation efficiency remains largely unexplored. Addressing these questions requires tools that can capture transcription factor behavior at high spatial and temporal resolution in live cells. Single-molecule imaging enables tracking of individual Pol II molecules during chromatin engagement, providing a powerful framework to dissect the kinetics of transcription initiation^9^. However, a comprehensive model linking TFIID and Pol II dynamics to initiation outcomes in vivo has yet to be established.

Here, we developed a live-cell single-molecule imaging approach that combines endogenously tagged Pol II and TFIID to dissect the kinetics of transcription initiation in human cells. By partitioning Pol II dwell times on chromatin and perturbing transcription with inhibitors, we classify binding events into distinct phases of initiation and early elongation, including promoter binding, proximal pausing, and productive gene body engagement. Our results reveal that transcription initiation is globally inefficient, highly heterogeneous between individual cells, and shaped by the dynamic order of pre-initiation complex assembly. These findings provide a new framework for understanding how kinetic and structural pathways jointly regulate transcription initiation in vivo.

## Results

### Generating an endogenously tagged human Pol II and TFIID cell line

To visualize the dynamics of transcription initiation in live cells, we generated a CRISPR knock-in U2OS cell line expressing SNAP-tagged RNA Polymerase II (Pol II) and Halo-tagged TFIID. The SNAP-tag was inserted at the N-terminus of the *POLR2A*, which encodes the large subunit of Pol II. To tag TFIID, we inserted a HaloTag at the C-terminus of *TAF1*, a gene encoding a TFIID specific subunit, via homology-directed repair (Fig. 1a, b; Supplementary Table 2). Dual knock-in lines were created by sequential editing with SNAP-Pol II was introduced first, followed by TAF1-HALO in the SNAP-Pol II background. Two rounds of fluorescence-activated cell sorting (FACS) were performed to first isolate cells containing either tag followed by a second round to enrich for homozygous integrations.

**Fig. 1:**
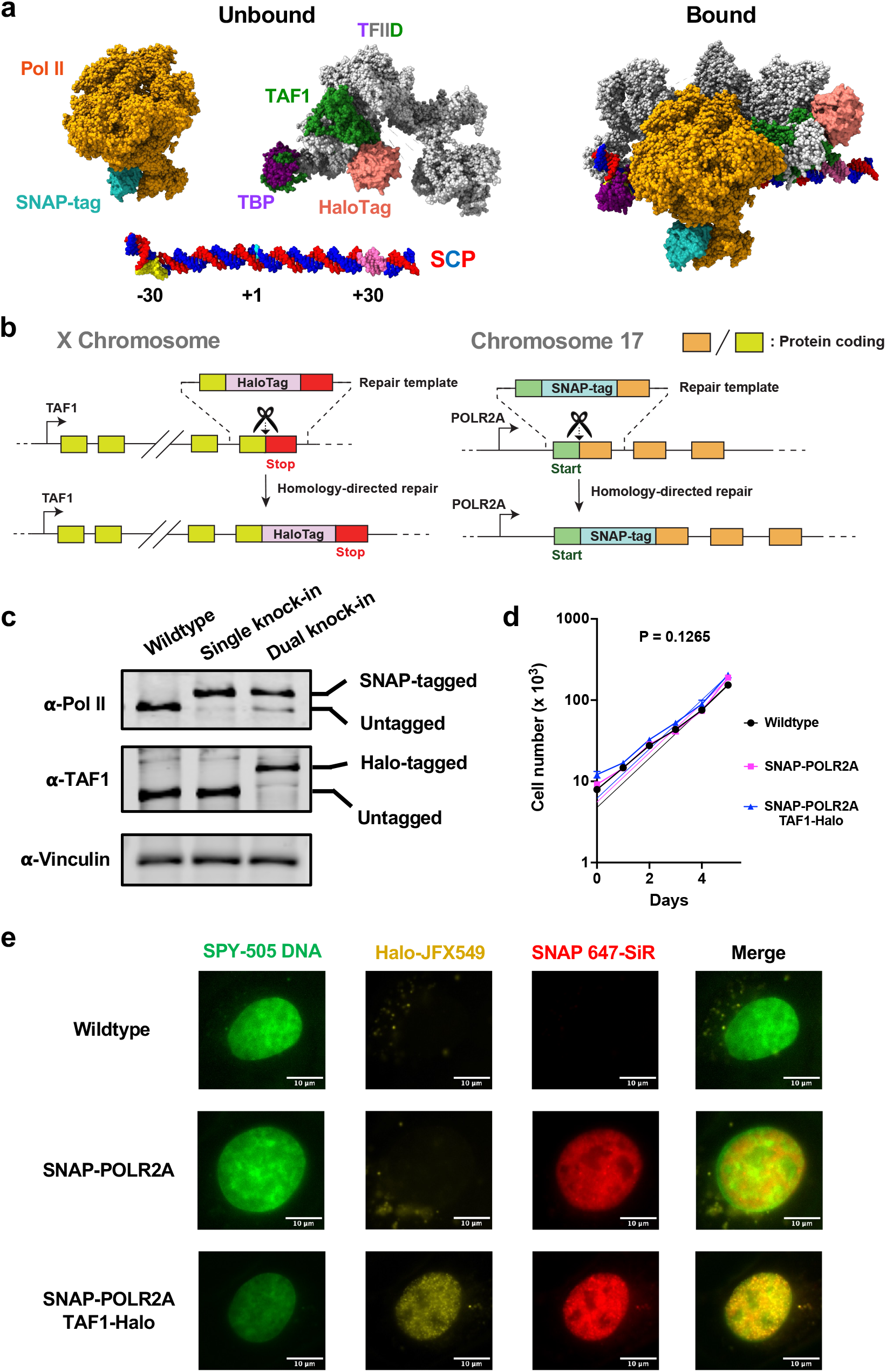
Establishing a CRISPR knock-in Pol II and TAF1 cell line. **a**, Schematic showing Pol II (PDB: 7NVS) and TAF1 (TFIID, PDB: 6MZL unbound; 6MZM bound) in an unbound (left) and bound (right) state at the super core promoter. SNAP (PDB: 6Y8P) and Halo (PDB: 6U32) tags were inserted into the N-terminus of the *POLR2A* gene and C-terminus of *TAF1*, respectively. Structural models were assembled using UCSF Chimera and are for illustrative purposes only; they are based on published PDB structures and do not represent an experimentally determined full complex. **b**, CRISPR-Cas9 strategy for dual knock-in. SNAP-tagged Pol II was generated first, followed by targeting the HaloTag into the C-terminus of TAF1 in the SNAP-Pol II background. **c**, Western blot analysis confirming expression of tagged proteins. Lysates from WT, single knock-in (SNAP-Pol II), and dual knock-in (SNAP-Pol II / TAF1-Halo) lines were probed with antibodies against Pol II, TAF1, and vinculin (loading control). Migration shifts indicate successful tagging. **d**, Growth curves showing that SNAP-Pol II and dual knock-in cells exhibit normal proliferation rates compared to WT. Data are plotted as mean ± SEM from triplicate wells. Curves were fit using a nonlinear Malthusian growth model (p value = 0.1265; null hypothesis: equal growth rate across all strains). **e**, Live-cell imaging of SNAP-Pol II and TAF1-Halo reveals nuclear localization of both tagged proteins. Cells were labeled with SPY505-DNA (Cytoskeleton), SNAP-Cell 647-SiR (NEB), and HaloTag JFX549 (Promega), and imaged using a single-frame 1-second exposure.

Immunoblotting confirmed expression of both tagged proteins with migration shifts of proteins relative to wild-type cells (Fig. 1c). Single and dual knock-in lines exhibited normal proliferation rates (Fig. 1d) and nuclear localization of the tagged proteins by live-cell imaging (Fig. 1e). Together, this system enables quantitative, endogenous analysis of Pol II and TFIID dynamics during transcription initiation in living cells.

### Single-molecule imaging reveals globally inefficient transcription initiation

During transcription initiation and early elongation, Pol II can dissociate at multiple regulatory checkpoints, including promoter instability, abortive initiation, and pausing near the +1 nucleosome^10-12^ (Fig. 2a). Only a small fraction of Pol II molecules are expected to transition into productive elongation by releasing from the proximal pause and entering the gene body. Indeed, previous work using fluorescence recovery after photobleaching (FRAP) at a single transgenic locus revealed that only ∼1% of alpha amanitin resistant Pol II molecules that bind the promoter ultimately engage in productive elongation, with the majority of Pol II dissociating during early initiation steps^7^. These results were further confirmed via global FRAP using endogenously tagged Pol II^8^. Together, these studies suggested that transcription initiation is highly inefficient. However, the conclusions drawn from these studies were based on a highly engineered locus-specific reporter system, ensemble measurements, and computational modeling.

**Fig. 2:**
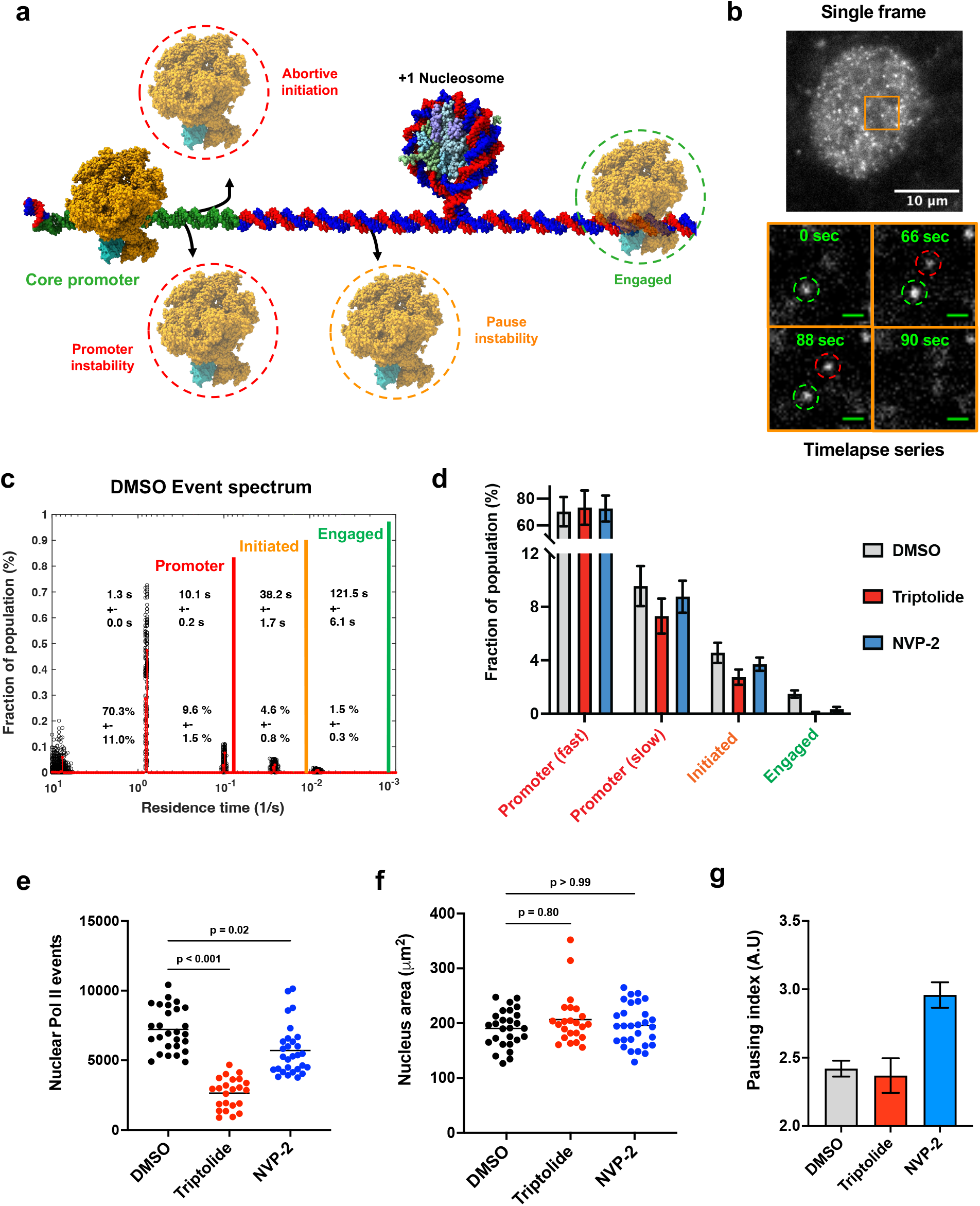
Single-molecule imaging reveals globally inefficient transcription initiation. **a**, Conceptual model illustrating kinetic loss of RNA polymerase II (Pol II) during distinct steps of transcription initiation. Most Pol II molecules dissociate at or shortly after promoter binding, due to promoter instability, abortive initiation, or promoter-proximal pause instability near the +1 nucleosome (PDB: 7Y8R; outlined in red). Only a small fraction of Pol II can successfully transition into productive elongation (outlined in green). **b**, Live-cell single-molecule imaging of SNAP-Pol II. Top: raw single frame showing sparse fluorescent spots. The orange box outlines a zoomed-in region shown below, where a time-lapse series (0–90 s) captures two individual Pol II binding events. One spot appears at 0 seconds (green outline) and a second spot appears at 66 seconds (red outline). Both spots persist through 88 seconds and disappear at 90 seconds, consistent with chromatin engagement and dissociation. **c**, GRID event spectrum of Pol II chromatin-binding events in DMSO-treated cells (1:1000, *n* = 27 cells). Four distinct kinetic populations were resolved, with mean residence times of ∼1.3, 10, 38, and 122 seconds. The least stable population (∼1.3 s) accounted for ∼70% of events, while the most stable population (∼122 s) comprised ∼1.5% of total events. Integration borders demarcating promoter, initiated, and engaged states were defined based on sensitivity to transcription inhibitors (see Supplementary Fig. 1b, c). Events with fitted lifetimes below the imaging interval (<1 second) were excluded from interpretation. **d**, Fraction of Pol II molecules in each of the four kinetic populations shown in panel c following treatment with 1 μM Triptolide (*n* = 23 cells) or 50 μM NVP-2 (*n* = 29 cells). Raw Pol II GRID plots for cells treated with these inhibitors can be found in Supplementary Fig. 1b and 1c. **e**, Total number of nuclear Pol II binding events per cell in DMSO (n = 27), Triptolide (n = 23), and NVP-2 (n = 29) conditions. Each point represents a measurement from a single nucleus. **f**, Total nuclear area imaged per cell in DMSO (n = 27), Triptolide (n = 23), and NVP-2 (n = 29) conditions. Each point represents a measurement from a single nucleus. **g**, Pausing index, calculated as the number of molecules with residence times between 40–90 s (paused) divided by those >90 s (elongated), aggregated across all cells per condition (DMSO, *n* = 27 cells; Triptolide, *n* = 23; NVP-2, *n* = 29 cells). Mean ± SEM is shown, with error estimated using Poisson statistics.

To determine whether this inefficiency is a global feature of endogenous transcription, we performed live-cell single-molecule imaging in U2OS cells expressing endogenous SNAP-tagged Pol II and Halo-tagged TFIID. We imaged cells under sparse-labeling conditions using a 500 millisecond exposure time to resolve individual chromatin-bound Pol II molecules^13^ (Fig. 2b). Fluorescent spots appeared and disappeared over time within the nucleus, reflecting dynamic Pol II engagement with chromatin. We extracted dwell times for ∼200,000 individual Pol II molecules across 27 nuclei and applied GRID (Genuine Rate Identification) analysis^14^ to identify discrete kinetic populations corresponding to Pol II dissociating during the phases of transcription initiation (Fig. 2a). GRID provided a superior fit to the dwell time distribution compared to conventional two- or three-exponential fit models (Supplementary Fig. 1a).

In DMSO-treated cells, GRID revealed four distinct binding populations with mean residence times of ∼1.3, 10, 38, and 122 seconds (Fig. 2c). The shortest-lived population (∼1.3 sec) accounted for ∼70% of binding events, while the most stable (∼122 sec) represented ∼1.5% of total events. To further characterize these kinetic populations, we treated cells with transcription inhibitors targeting different steps of the initiation process. Triptolide, an XPB inhibitor that blocks promoter opening and promoter escape^15,16^, substantially reduced the percentage of Pol II occupying both intermediate (∼38 sec) and stable fractions (∼122 sec) (Fig. 2d; Supplementary Fig 1b). Therefore, we have labeled the intermediate population as Pol II in an ‘initiated’ state, meaning it escaped the promoter but did not transition into a stably engaged state associated with productive elongation into the gene body (Fig. 2c). In contrast, treatment with NVP-2, a specific CDK9 inhibitor that blocks pause release^17^, selectively reduced Pol II’s occupancy of only the most stable fraction, which we define as Pol II that has reached an “engaged” state and transcribed into the gene body (Fig. 2c, d; Supplementary Fig 1c). These trends were consistent across two independent imaging sessions (Supplementary Fig. 1d).

To further understand how drug treatment impacts Pol II binding to the genome, we analyzed individual nuclei from drug-treated U2OS cells and quantified the total number of Pol II binding events per nucleus. Both Triptolide and NVP-2 treatments led to a reduction in nuclear Pol II events compared to DMSO (Fig. 2e). Nuclear area did not significantly differ between conditions (Fig. 2f), ruling out nuclear size as a confounding factor (see Supplementary Fig. 1e, f for replicate data). The reduction in Pol II signal following Triptolide is consistent with previous reports showing that Triptolide induces proteasome-mediated degradation of Pol II’s largest subunit (RPB1) in a time and dose dependent manner^18^. Our treatment conditions (1 µM for 2 hours) exceed the concentrations and durations previously shown to deplete Pol II protein levels, suggesting that the observed loss in nuclear binding events likely reflects both transcriptional shutdown and Pol II degradation. For NVP-2, the more subtle drop in Pol II binding events likely reflects fewer molecules detected during the late stages of initiation due to impaired pause release and reduced transcriptional burst frequency^19^.

### Kinetic partitioning of Pol II binding events recapitulates global proximal pausing

Based on prior FRAP studies, promoter-proximal pausing is estimated to occur ∼40 seconds after Pol II engages the promoter^8^. In our GRID analysis, we observed a distinct population of Pol II molecules with residence times in this range, which failed to transition into an engaged state following CDK9 inhibition with NVP-2. This supports the idea that Pol II pauses ∼40 seconds on average after binding and that CDK9 is required for pause release. Previous estimates using genomic methods placed the median promoter-proximal pause duration at ∼60-70 seconds^20^. Therefore, we conservatively defined Pol II elongation into the gene body as molecules with dwell times exceeding 90 seconds. Pol II molecules exceeding this threshold are also sensitive to CDK9 inhibition and approach the photobleaching limit (∼128 sec, Supplementary Fig. 1e), supporting their classification as elongating into the gene body.

Using this classification, we defined a global cellular pausing index as the ratio of the number of Pol II molecules with residence times between 40–90 seconds (paused) to those exceeding 90 seconds (gene body). CDK9 inhibition is known to increase the pausing index by impairing pause release^21,22^. Consistent with this, NVP-2 treatment increased the pausing index relative to control (Fig. 2g; Supplementary Fig. 1h). In contrast, Triptolide, which blocks promoter escape, reduced both paused and gene body fractions proportionally, resulting in no net change in the pausing index.

Together, these results suggest that Pol II binding events partition into four kinetically distinct states. GRID fitting indicated two states corresponding to unstable promoter interactions, one representing an intermediate initiated state, and a rare, long-lived state (∼122 sec) likely associated with productive elongation into the gene body. Independent of fitting methods, Pol II binding events can further be partitioned based upon dwell time into a paused (40-90 sec) and a gene body (>90 sec) state, which can be selectively targeted by CDK9 inhibition.

### Single-cell analysis reveals heterogeneity associated with Pol II entering into productive elongation

We define transcription efficiency as the fraction of Pol II molecules that reach the most stable kinetic population (Pop 4), representing long-lived Pol II that has transcribed into the gene body. To assess whether this efficiency varies between individual cells, we performed single-cell GRID analysis. In DMSO treated cells, we observed substantial heterogeneity in the fraction of Pol II molecules reaching the engaged state, defined by long residence times near or beyond the photobleach limit. For example, one cell exhibited ∼2.8% of molecules in Pop 4 with a residence time of ∼98 seconds (Fig. 3a), while another showed only ∼0.4% at ∼152 seconds (Fig. 3b).

**Fig. 3:**
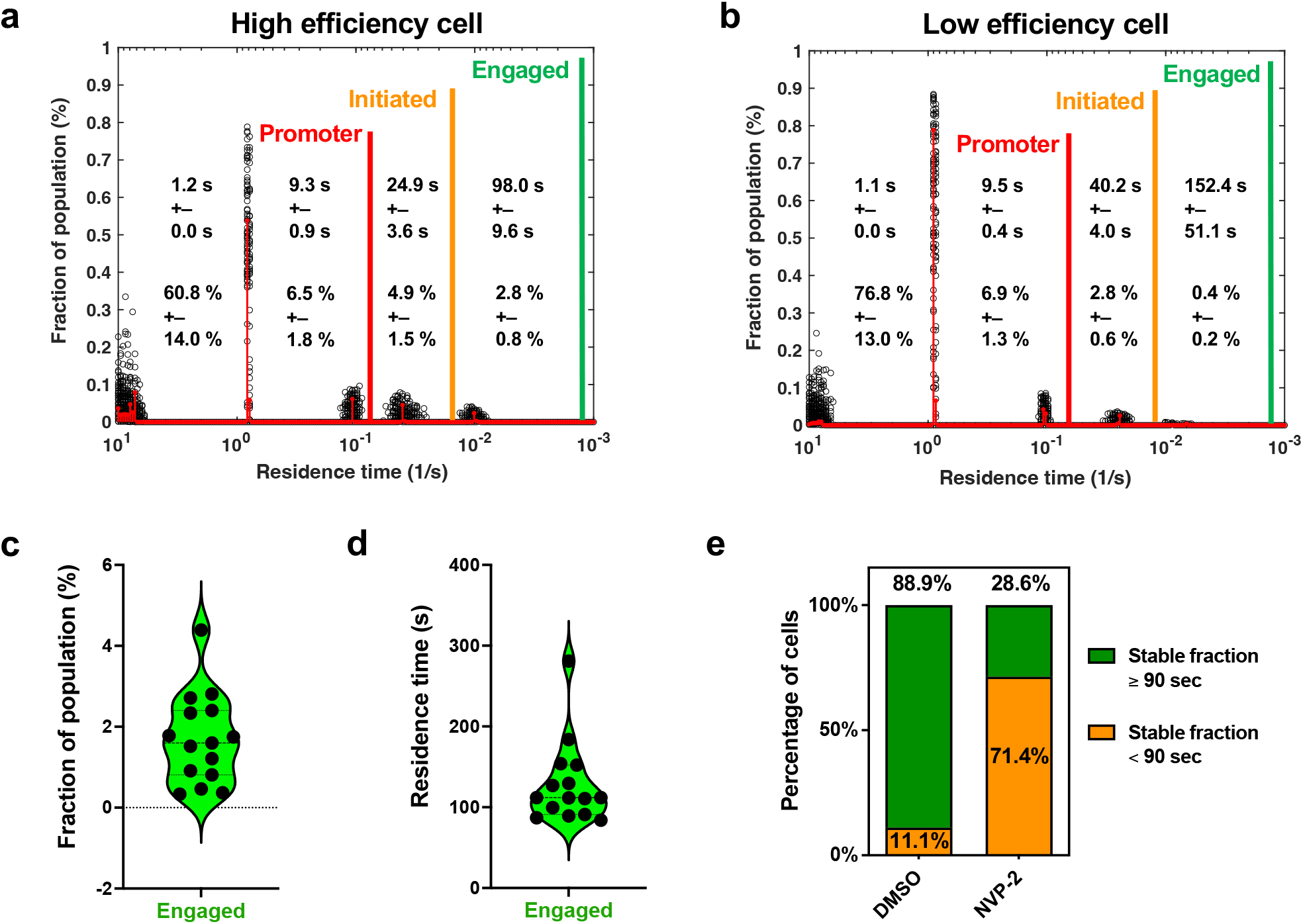
Single-cell analysis reveals heterogeneity in transcription efficiency. **a**, GRID event spectrum from a DMSO-treated nucleus exhibiting high transcription efficiency. This cell shows a higher fraction (∼2.8%) of Pol II molecules reaching the engaged state, with a residence time within error of the photobleach limit (∼106 s), indicating productive elongation. **b**, GRID event spectrum from a DMSO-treated nucleus exhibiting low transcription efficiency. This cell shows a lower fraction (∼0.4%) of Pol II molecules reaching the engaged state, with a residence time within error of the photobleach limit (∼148 s). **c**, Distribution of the fraction of Pol II molecules in the engaged state (Pop 4) across individual DMSO-treated cells (n = 27). Right: distribution of corresponding residence times for Pop 4. Only cells with GRID fits accounting for >94% of events were included. **d**, Distribution of the residence times for Pol II molecules in the engaged state (Pop 4) across individual DMSO-treated cells (n = 27). Only cells with GRID fits accounting for >94% of events were included. **e**, Percentage of cells in DMSO and NVP-2 treated conditions with the most stable population (Pop 4) having a residence time equal to or exceeding 90 seconds (stably engaged state). This was used as a proxy for productive elongation.

Across all DMSO-treated nuclei, the fraction of molecules in Pop 4 ranged from 0.3% to 4.4%, with a broad distribution of residence times (Fig. 3c, d). However, a number of cells only contained 3 populations consisting of short and intermediate Pol II binding molecules and therefore lacked engaged Pol II. To assess how many cells exhibited engaged Pol II, we calculated the percentage of cells in each condition with residence times equal to or exceeding 90 seconds in their most stable fraction. Almost all DMSO-treated cells met this threshold (∼89%), while this fraction dropped sharply in NVP-2–treated cells (∼29%, Fig. 3e), consistent with a loss of productive elongation following transcription inhibition. Together, these results demonstrate that transcription efficiency is highly heterogeneous at the single-cell level.

### TFIID-associated Pol II has distinct transcriptional efficiencies relative to global Pol II binding events

To characterize the dynamic assembly of functional Pol II pre-initiation complexes in live cells, we utilized multi-color imaging data combined with our Single-Molecule Analysis of Colocalization Kinetics (SMACK) pipeline. Two-dimensional projections revealed widespread colocalization of Pol II and TFIID signals within the nucleus (Fig. 4a, top left). A representative time-lapse series illustrated a colocalization event where TFIID appears prior to Pol II with Pol II persisting after TFIID dissociation. This order of assembly pattern is similar to canonical transcription initiation from a TFIID directed pre-initiation complex, based upon in vitro biochemistry and structural studies^3,5,23^ (Fig. 4a, bottom schematic). To further characterize detected assembly events, we measured the distance between temporally and spatially co-localized Pol II and TFIID binding trajectories. The majority of colocalization events (∼72%) occurred at a distance of less than a single pixel (∼110 nm) with a median distance of ∼79 nm (Fig. 4b). This median co-localization distance is equivalent to roughly 232 bp of linear B-form DNA.

**Fig. 4:**
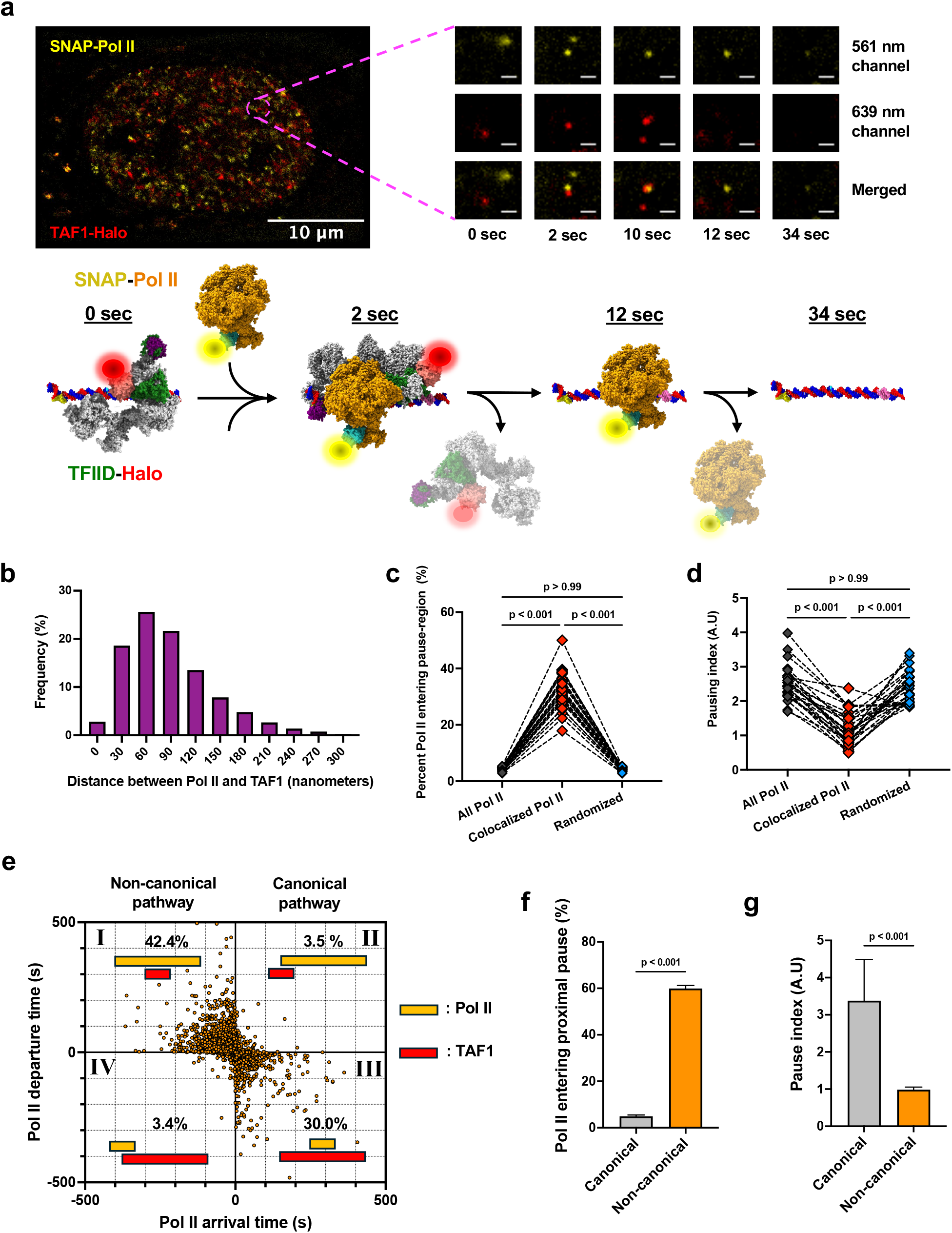
Non-canonical colocalization of Pol II and TAF1 drives higher transcriptional efficiency and reduced pausing. **a**, Two-dimensional projection of a U2OS nucleus showing SNAP-tagged Pol II (yellow) and Halo-tagged TAF1 (red). A zoomed-in region (right) shows a representative colocalization event across five time points (0, 2, 10, 12, and 34 s) in the 561 nm (Pol II), 639 nm (TAF1), and merged channels. Scalebar indicates a distance of 1 micron. A schematic illustrating the inferred order of binding from selected time points is shown below. **b**, Histogram showing the average distance between TAF1 and Pol II trajectories during colocalization events. **c**, Percentage of Pol II molecules entering the pause-region (Pol II residence time ≥40 s) for all Pol II, colocalized Pol II, and randomized controls. Single-cell paired analysis bootstrapped 1,000 times per cell. Statistical significance assessed using a Friedman test. **d**, Pausing index, calculated as the number of Pol II molecules with residence times between 40–90 seconds divided by those >90 seconds, using the same groups and bootstrapping strategy as in (c). Statistical significance assessed using a Friedman test. **e**, Quadrant analysis of colocalization dynamics, with Pol II arrival relative to TAF1 on the X-axis and departure on the Y-axis. Percentages indicate the fraction of events in each quadrant. Quadrants I and IV (upper left, bottom left, respectively) represent non-canonical dynamics (Pol II arrives first); quadrants II and III (upper right, bottom right, respectively) represent canonical dynamics (TAF1 arrives first or simultaneously with Pol II). **f**, Percentage of Pol II molecules that reach proximal-pause for canonical and non-canonical colocalization events, computed by bootstrapping 80% of events over 1,000 iterations. Bar plots in show means ± bootstrapped standard deviations. **g**, Pausing index for canonical versus non-canonical colocalization events using the same bootstrapping strategy as in (f). Bar plots in show means ± bootstrapped standard deviations.

Given that TFIID binding drives Pol II recruitment, we wanted to understand how the presence of TFIID impacts the ability of Pol II to efficiently escape the promoter and enter the pausing region (∼+30 to +80bp) of genes. Therefore, we compared the fraction of Pol II molecules with residence times ≥40 seconds (based on our GRID analysis of productive initiation events) for all Pol II molecules, TFIID colocalized Pol II molecules, and randomized controls. The percentage of Pol II entering the pause region was significantly higher for TFIID colocalized Pol II molecules relative to all Pol II and randomized controls (Fig. 4c). Entry of colocalized Pol II into the pause region was also significantly decreased after treatment with Triptolide and NVP-2 (Supplemental Fig. 3a). TFIID colocalized Pol II molecules also exhibited a lower pausing index, defined as the ratio of molecules with residence times between 40–90 seconds (paused) to those >90 seconds (elongating into gene body, Fig. 4d). These results suggest that Pol II colocalization with TFIID is associated with more efficient pause release compared to global Pol II.

### Transcriptional outcomes are dependent on TFIID and Pol II assembly pathways

To further dissect the dynamics of TFIID and Pol II assembly, we performed quadrant analysis of the arrival and departure times of Pol II with respect to TFIID (Fig. 4e). Only 3.5% of events exhibited a canonical assembly pathway, where TFIID arrives first and Pol II leaves last (quadrant II, top right). Of note, 30% of the colocalization events arise from TFIID arriving first and Pol II dissociating before TFIID falls off the genome. This canonical assembly pathway is structurally incompatible with Pol II escaping past ∼+13-18bp as seen in a cryo-EM study of elongating Pol II^24^. These events likely reflect a large degree of abortive initiation events associated with TFIID driven pre-initiation complex formation and contribute to substantial transcription initiation inefficiency. Surprisingly, a majority of colocalization events (42.4%) fell into quadrant I (top left), where Pol II arrives first and leaves last relative to TFIID, potentially representing a non-canonical pathway. Alternatively, these co-localization events could arise from Pol II being loaded onto the promoter in a canonical manner via a “dark” or unlabeled TFIID.

Given the dominance of non-canonical colocalization, we compared the efficiency of pause entry (e.g. Pol II >40”) and the pausing index between events displaying canonical and non-canonical assembly pathways. Non-canonical colocalization assembly events displayed a higher pause entry efficiency (Fig. 4f) and a lower pausing index (Fig. 4g) relative to canonical assembly events. The fact that we see these differences between the non-canonical and canonical pathways argues against non-canonical assembly arising from recruitment of Pol II via a “dark” TFIID binding event. Furthermore, these results suggest that colocalization events where TFIID binds before Pol II lead less efficient pause entry and a stronger pausing index for Pol II molecules that do escape the promoter and enter the pausing region (Fig. 5). Correspondingly, for events where TFIID binds after Pol II has been loaded and escaped the promoter, Pol II is associated with more efficient entry into the pause region and more efficient pause release. This potentially reflects genes with more accessible chromatin and increased transcriptional permissiveness leading to Pol II convoy formation and transcriptional bursting. Together, these findings reveal that the local dynamics and assembly pathways of Pol II and TFIID interactions correlate with distinct transcriptional outcomes, highlighting nuclear subdomains that favor rapid initiation and pause release.

**Fig. 5:**
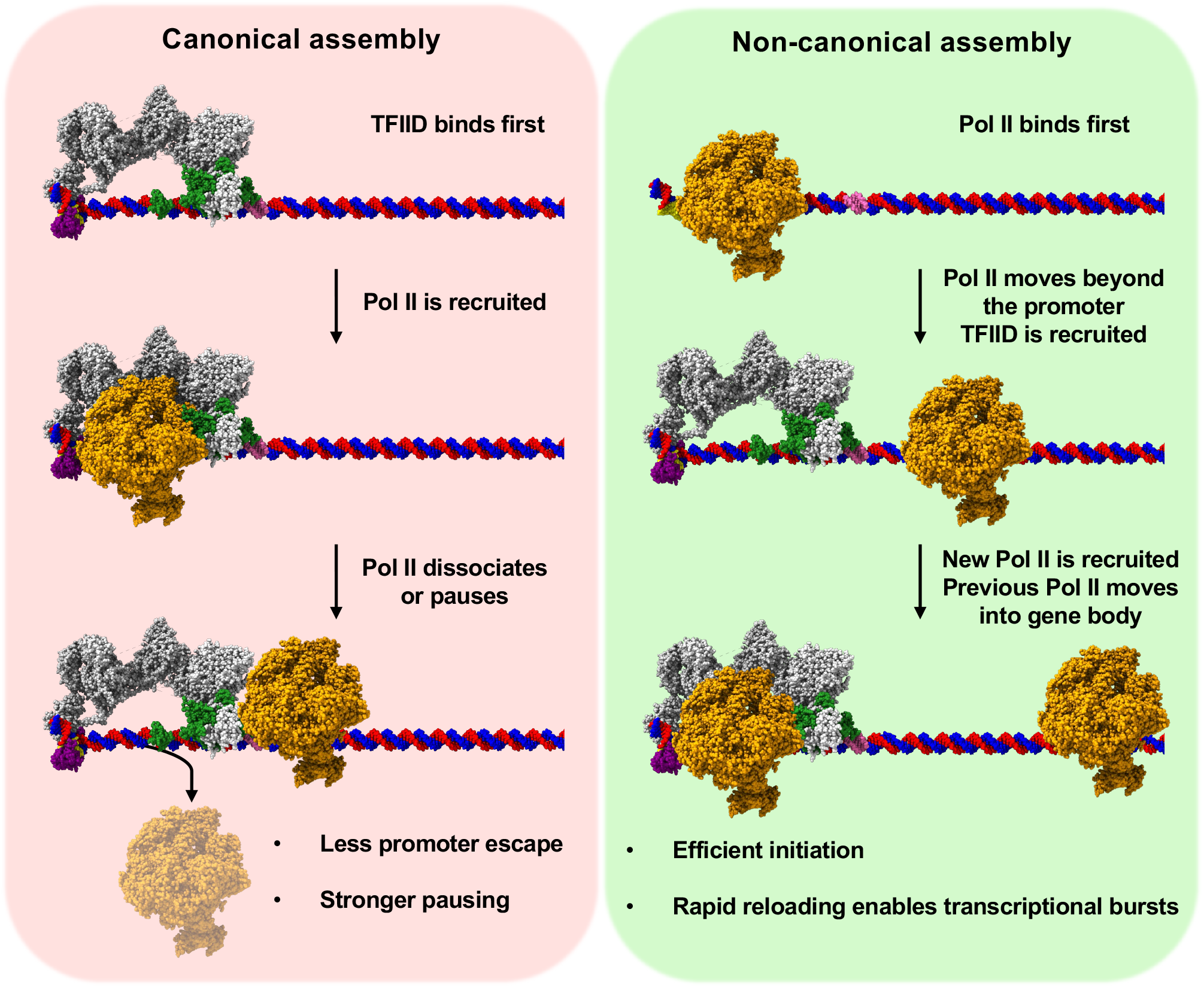
Schematic model and features of canonical versus non-canonical assembly of pre-initiation complexes. In the canonical assembly (left), TFIID arrives first followed by Pol II. This type of pre-initiation complex frequently leads to abortive initiation or for Pol II that escapes the promoter, a strongly paused complex. The non-canonical assembly (right) represents a situation where Pol II is loaded onto the gene by an unknown (not TFIID) core promoter binding complex. At some point after Pol II initiation, TFIID arrives and likely helps recruit a trailing Pol II during a transcriptional burst. The initial Pol II from a non-canonical assembly is highly efficient at escaping the promoter with a low propensity to pause.

## Discussion

There are now two orthogonal imaging approaches (FRAP and single molecule tracking) to probe functional Pol II dynamics in live cells. Live cell single molecule imaging allows us to quantify these different Pol II binding states which yield similar values yet are independent of complex modeling of ensemble FRAP recovery curves. Using transcriptional inhibitors targeting various stages of initiation and early elongation, we can now begin to determine the percentage of Pol II entering in these different phases and their kinetic properties. Importantly, both approaches indicate that functional transcription initiation is extremely inefficient inside live cells, with approximately 1% of Pol II binding events leading to a long-lived state that is sensitive to pause release inhibition^7,8^. Inefficient transcription initiation is likely due a number of quality control stages, including abortive initiation, 5’ end capping, engagement with the +1 nucleosome, and proximal pausing. This stringent level of control ensures that Pol II assembles regulators necessary for capping, elongation and termination before committing to elongation into the gene body. This inefficiency suggests that most genes exist in a default “off” or idling state, awaiting regulatory signals that transiently enhance transcription efficiency.

Our Pol II dwell times measured from live cell imaging and Pol II kinetics measured from genomic assays are remarkably similar^12^. This provides a unified kinetic pathway suggesting that Pol II enters into the gene body rather slowly (∼60-90 sec) after landing on the promoter and proximally paused Pol II is overall dynamic lasting in the range 45-180 seconds on average even after inhibition of CDK9^25^. By classifying these different transcriptional states based upon our single molecule derived Pol II dwell times (paused, 40-90 sec and gene body, >90 sec), we can recapitulate a global cellular pausing index that is sensitive to CDK9 inhibition.

Due to the sensitivity of our imaging assay, we obtain a pausing index for individual cells which varies considerably across our population of cells, ranging from 1.7 to 4.0 (Fig. 4d). We see a similar high variability in the percentage of Pol II molecules that are long-lived and likely functional engaged or elongated into the gene body (Fig. 3c). This large spread in functional engagement and pausing index suggests different cell states where regulation of proximal pause release differs. We can’t yet determine the source of the cellular variability in regulation of Pol II engagement; however, we speculate that regulation of Pol II proximal pausing will differ depending on the phase of cell cycle. For example, it is well known that transcription is suppressed during early S phase, when DNA Polymerase is replicating euchromatic regions^26^. In such a scenario, it might be advantageous for the pausing index to be high with less Pol II engaged in the gene body. In addition, cells approaching mitosis might have a high global pausing index to accommodate known mitotic bookmarking of Pol II^27,28^. Correspondingly, global pause index might be low in cells in G1 or early G2 where transcription rates are predicted to be higher than in S phase or mitosis.

Live cell multi-color single molecule imaging is a powerful tool to assess functional assembly pathways inside cells^29^. By simultaneously imaging TFIID and Pol II assembly, we find a number of important features to classify different Pol II binding states. Compared to visualizing global Pol II binding alone, colocalization of Pol II with TFIID leads to a dramatic increase (from ∼4% global to 18-50% colocalized) in the efficiency of Pol II lasting long enough to enter into a paused state. This is due to the fact that global Pol II binding events likely represent a significant amount of Pol II molecules that are searching but not yet finding promoter regions. The observation that Pol II pausing behavior shifts depending on the spatial and temporal context of TFIID binding supports the idea that our colocalization measurements capture functional interactions. This is further reinforced by the finding that Pol II colocalization with TFIID is associated with a lower pausing index relative to global Pol II binding events.

It was surprising to see that Pol II associated with TFIID had a lower pausing index, given that a prior study associated TFIID with an increase in Pol II proximal pausing^30^. This is reconciled by temporal analysis revealing that colocalization events could be broken down into a canonical pathway, where TFIID binds before Pol II, and a non-canonical pathway, where TFIID binds after Pol II. The canonical pathway has a pausing index that is roughly equivalent to global Pol II (3.3 canonical vs. 2.5 for global, Fig. 4d, g). In contrast, the non-canonical pathway where Pol II arrives first has a much lower pause index (∼1, Fig. 4g).

Surprisingly, the canonical pathway, where TFIID dissociates before Pol II, is exceedingly rare (∼3.5% in Quadrant II, Fig. 4e). Instead, most canonical events involve Pol II binding after TFIID and dissociating before TFIID release (∼30% in Quadrant III, Fig. 4e). This order of assembly is consistent with both structural models and our kinetic data, which show that Pol II frequently dissociates before successfully escaping the promoter. Accordingly, the low pausing index observed for colocalized Pol II likely reflects the scarcity of long-lived, canonically assembled molecules relative to the more frequent, non-canonical Pol II events.

The large difference in pausing behavior between canonical and non-canonical events may reflect recruitment by a factor or complex distinct from TFIID, with TFIID instead stabilizing the promoter for subsequent Pol II binding. Supporting this possibility, studies in budding yeast have shown that SAGA is required for TFIID-mediated transcription on a subset of genes^31^. In such a scenario, SAGA recruits an initially elongation-prone Pol II, with TFIID contributing to the recruitment of additional polymerases during transcriptional bursting. In this model, de novo TFIID binding may serve as a stochastic brake that introduces promoter-proximal pausing, enabling more precise regulation of transcriptional output.

Overall, our dynamic, multi-color, live-cell single-molecule imaging approach offers key advantages over ensemble and genomic methods, including the ability to capture cellular heterogeneity and directly extract kinetic rates for distinct steps in transcription. These capabilities provide new insights into how regulatory factors influence functional Pol II engagement in vivo. Moving forward, we envision applying this approach to additional components of the pre-initiation complex and chromatin-modifying factors. However, current limitations prevent the direct measurement of gene-specific kinetic parameters and pausing indices. Addressing this challenge will require the development of methods capable of tracking multiple regulatory factors at defined genomic loci in real time within single cells.

## Methods

### Cell line generation using CRISPR/Cas9 genome editing

Human osteosarcoma (U-2 OS) cells (ATCC HTB-96) were cultured at 37°C with 5% CO_2_ in high-glucose DMEM (Gibco, 10569044) supplemented with 10% fetal bovine serum (FBS, vol/vol) and 1% penicillin-streptomycin (500 U/mL). Cells were passaged at ∼70% confluency in 10 cm tissue culture dishes.

Guide RNAs (gRNAs) were prepared by mixing 3.6 μL of 200 μM tracrRNA (IDT, 1072532) with 3.6 μL of 200 μM gene-specific crRNA (Supplementary Table 1) in nuclease-free duplex buffer (IDT, 11010301). The mixture was incubated at 95°C for 5 minutes and allowed to cool to room temperature for 20 minutes to facilitate crRNA-tracrRNA duplex formation. The gRNA targeted a region of the N-terminus of human *POLR2A* spanning start codon on exon 1.

Ribonucleoprotein (RNP) complexes were formed by combining 7.2 μL of gRNA, 4 μL of HiFi Cas9 Nuclease (IDT, 1081060), and 0.8 μL of PBS at room temperature for 20 minutes.

Homology-directed repair (HDR) templates were generated from PCR-amplified double-stranded gBlock HiFi gene fragments (IDT, for SNAP-POLR2A) or clonal genes (Twist Bioscience, for TAF1-Halo) containing either the SNAP or HaloTag flanked by sequence specific homology arms (Supplementary table 2, 3). The amplified double-stranded DNA sequences were converted to single-stranded DNA using the Guide-it long ssDNA kit (Takara, 632666).

Approximately 1 million cells were resuspended in 100 μL of nucleofection solution containing 20 μL of ssDNA (∼5 μg), 10 μL of the RNP mix, and 70 μL of Nucleofector solution V + Supplement I (mixed at a 4.5:1 ratio; Lonza, VCA-1003). The resuspended cells were transferred to an aluminum cuvette (Lonza) and electroporated using the Amaxa Nucleofector II with the X-001 program. Immediately after electroporation, 300 μL of pre-warmed culture media was added to the cuvette and gently mixed before plating in a 6-well culture plate containing 1.6 mL of pre-warmed culture media (DMEM + 10% FBS + 1% penicillin-streptomycin) supplemented with 3 μL of 690 μM Alt-R HDR Enhancer V2 (∼1 μM final; IDT, 10007910). Cells were incubated at 37°C with 5% CO_2_ for 24 hours before being washed with PBS and adding fresh culture media. Once cells reached ∼70-80% confluency, they were expanded into progressively larger vessels and sorted by fluorescence activated cell sorting (FACS) based on SNAP-Cell 647-SiR dye fluorescence (NEB, S9102S).

### Generating dual SNAP- and Halo-tag knock-in lines

To generate the dual knock-in SNAP-POLR2A/TAF1-Halo cell lines, validated SNAP-POLR2A cells were re-targeted using a gRNA specific to the C-terminus of human TAF1. RNP complexes were formed as described above, using a TAF1-specific crRNA and a HaloTag repair template obtained as a clonal gene fragment (Twist Bioscience) with homology arms flanking the stop codon on exon 39. Single-stranded DNA was generated as previously described.

Approximately 1 million SNAP-POLR2A cells were electroporated with the RNP complex and TAF1-Halo ssDNA repair template using the same nucleofection conditions. Dual-labeled cells were identified using Halo JFX549 dye (Janelia, gift from Luke Lavis) and sorted via FACS. A second round of FACS was performed to enrich for a stable, high-penetrance polyclonal population. These dual knock-in strains were expanded and used for all subsequent experiments.

### Cell line validation assays

For Western blotting, whole-cell extracts were harvested from ∼70% confluent 15 cm tissue culture dishes using a cell scraper on ice and lysed in RIPA buffer supplemented with protease and phosphatase inhibitors (Thermo, 78442). Lysates were sonicated using a Diagenode Bioruptor (30 seconds on/off, high power, 5 minutes) and clarified by high-speed centrifugation at 18,000 x *g* for 15 minutes at 4°C. Protein concentration was determined using a detergent-compatible Bradford assay (Thermo, 23246). Equal amounts of total protein (30 μg) were resolved on 6% SDS-PAGE gels and transferred to Amersham Protran 0.45 μm nitrocellulose membranes (Cytvia, 10600003) using a wet transfer.

Membranes were blocked in 5% non-fat dry milk (Bio-Rad, 1706404) in 1x TBS for 1 hour at room temperature and incubated overnight at 4°C with the following primary antibodies: anti-Pol II (A-10) mouse monoclonal (Santa Cruz, sc-17798, 1:1000 dilution), anti-TAF1 (D6J8B) rabbit monoclonal (Cell Signaling Technologies, 12781, 1:1000 dilution), and anti-Vinculin (E1E9V) rabbit monoclonal (Cell Signaling Technologies, 13901, 1:1000 dilution) as a loading control. Blots were incubated with Alexa Fluor-conjugated goat anti-rabbit secondary antibody (Invitrogen, A32735, 1:20,000) or IRDye-conjugated goat anti-mouse secondary antibody (LI-COR, 926-32210, 1:20,000) and imaged using an Odyssey CLx imaging system.

To assess whether the SNAP- or Halo-tagged cell lines maintained normal growth rates, 10,000 cells were seeded in a 6-well plate. Cell growth was quantified every ∼24 hours for 5 days using manual counting with a grid reticle. Each cell line was quantified in triplicate, and data were plotted as mean ± SEM.

For localization, 7×10^5^ cells were seeded on 35 mm imaging dishes (MATTEK, P35G-1.5-14-C) 24 hours before imaging. Cells were labeled with SPY505-DNA (Cytoskeleton, CY-SC101, 1:1000 in DMSO, 1 hour), SNAP-Cell 647-SiR (100 nM, 30 minutes), and HaloTag JFX549 (250 pM, 30 minutes) at 37°C. After labeling, cells were washed with pre-warmed PBS before adding imaging media, which consists of Leibovitz’s L-15 (Fisher, 21083027) + 10% FBS. A single, 1 second exposure image was captured on a customized inverted Nikon Eclipse Ti microscope equipped with a 100x oil-immersion objective (NA 1.49, Nikon). Fluorophores were excited using 488 nm, 561 nm, and 639 nm lasers, and images were captured using a Prime 95B sCMOS camera.

### Single-molecule imaging

Live-cell single-molecule imaging was performed on dual knock-in U2OS cells expressing SNAP-tagged Pol II and Halo-tagged TAF1. Cells were labeled with 6 nM SNAP-Cell TMR-Star (NEB, S9105S) and 5 nM HaloTag JF646 ligand (Promega, HT1060) for 20 minutes at 37°C in pre-warmed DMEM. After labeling, cells were washed with complete culture media and pre-warmed PBS. Cells were then incubated for 2 hours at 37°C in 2 mL of Leibovitz’s L-15 medium (Thermo Fisher, 21083027) supplemented with 10% FBS prior to imaging. During this incubation, cells were treated with 2 μL of DMSO, 1 mM Triptolide, or 50 μM NVP-2.

Single-molecule imaging was performed on a customized Nikon Eclipse Ti inverted microscope equipped with a 100× oil-immersion objective (NA 1.49, Nikon), a Prime 95B sCMOS camera (Teledyne Photometrics), and 561 nm and 640 nm lasers for excitation. Images were acquired every 2 seconds for ∼16 minutes (with a 1.5-second dark time) under highly sparse labeling conditions to ensure single-molecule resolution. Although both channels were imaged, only the SNAP-Pol II (TMR) channel was analyzed for Figure 2 and 3.

Raw image stacks were processed using the STRAP and SMACK pipelines for background subtraction, spot tracking, colocalization analysis, and kinetic quantification. Both pipelines are available via the repositories listed in the Code availability section.

### GRID analysis

Dwell time distributions were analyzed using GRID (Genuine Rate Identification), as previously described (ref: Gebhardt lab). The event spectrum was used to determine the fraction of total Pol II binding events within each kinetic population during the imaging window. Events with fitted lifetimes below the imaging interval (<1 second) were excluded from biological interpretation. For single-cell GRID analysis, only cells with fits accounting for >94% of events were included. Aggregated GRID data were resampled 100 times using 80% of the data; single-cell GRID was resampled 100 times using 90% of each cell’s data.

## Author contributions

**NH:** Designed and performed all experiments, including CRISPR knock-in generation, single-molecule imaging, drug treatments, and quantitative analyses. Developed the STRAP and SMACK processing and analysis pipelines. Prepared all figures and wrote the manuscript. **RAC:** Supervised the project, helped develop the SMACK pipeline, guided experimental design and data interpretation, and wrote manuscript.

## Acknowledgements

We thank the Albert Einstein College of Medicine Flow Cytometry Core for assistance with cell sorting. We are grateful to the laboratory of Robert Singer for access to the nucleofector, which enabled CRISPR knock-in experiments. We also acknowledge Shuhao Liou for early work related to transcription inhibitor treatments in exogenous Pol II systems. This work was supported by grants from the National Institutes of Health (NIH), including NIGMS R01GM126045 and the Chan Zuckerberg Initiative Metabolism Network (MET-0000000459) awarded to R.A.C. and by the National Science Foundation Graduate Research Fellowship under Grant No. FAIN-2437848 awarded to N.H.

## Code availability

The STRAP (Single-molecule TRacking And Processing) and SMACK (Single-Molecule Analysis of Colocalization Kinetics) pipelines used in this study are available at https://github.com/codedbynayem/STRAP and https://github.com/codedbynayem/SMACK. STRAP integrates Fiji, Python, and MATLAB modules for preprocessing and tracking single-molecule imaging data. SMACK analyzes colocalization dynamics and computes kinetic parameters from time-resolved dual-channel data. Both pipelines are available under the GNU General Public License v3.0.

## Notes

### Competing Interest Statement

The authors have declared no competing interest.

