## Supplemental File for "Dynamic transcription pre-initiation complex assembly governs initiation efficiency"

### Supplementary information

| | Sequence (5’ to 3’) |
|---|---|
| *POLR2A* crRNA | GCCACCCCCGTGCATGGCGG |
| *TAF1* crRNA | GACTTGGACTCTGATGAATG |

**Supplementary Table 1: crRNA sequences targeting human *POLR2A* and *TAF1* genes.**

| SNAP-*POLR2A* HDR Repair template sequence (5’ to 3’) |
| --- |
| GCTCAGAAGCGCCGAGAGCGCGGCCGGGACGGTTGGAGAAGAAGGCGGCTCCCGGAAGGGGGAGAGACAAACTGC<br>CGTAACCTCTGCCGTTCAGGAACCCGGTTACTTATTTATTCGTTACCCTTTTTCTTCTTCCTCCCCCAAAAACCT<br>TTTCCTTTTCCCTTCTTTTTTTTTCCTTTTTGGGAGCTGAAAAATTTCCGGTAAGGGAAAGAAGGGCTCCTTTCG<br>CTCCTTATTTCCCCGCCTCCTTCCCTCCCCCACCTTCCCCTCCTCCGGCTTTTTCCTCCCAACTCGGGGAGGTCC<br>TTCCCGGTGGCCGCCCTGACGAGGTCTGAGCACCTAGGCGGAGGCGGCGCAGGCTTTTTGTAGTGAGGTTTGCGC<br>CTGCGCAGCGCGCCTGCCTCCGCC**ATG**TGGAGCCACCCGCAGTTCGAAAAAACCGGT**GACAAAGACTGCGAAATG**<br>**AAGCGCACCACCCTGGATAGCCCTCTGGGCAAGCTGGAACTGTCTGGGTGCGAACAGGGCCTGCACCGTATCATC**<br>**TTCCTGGGCAAAGGAACATCTGCCGCCGACGCCGTGGAAGTGCCTGCCCCAGCCGCCGTGCTGGGCGGACCAGAG**<br>**CCACTGATGCAGGCCACCGCCTGGCTCAACGCCTACTTTCACCAGCCTGAGGCCATCGAGGAGTTCCCTGTGCCA**<br>**GCCCTGCACCACCCAGTGTTCCAGCAGGAGAGCTTTACCCGCCAGGTGCTGTGGAAACTGCTGAAAGTGGTGAAG**<br>**TTCGGAGAGGTCATCAGCTACAGCCACCTGGCCGCCCTGGCCGGCAATCCCGCCGCCACCGCCGCCGTGAAAACC**<br>**GCCCTGAGCGGAAATCCCGTGCCCATTCTGATCCCCTGCCACCGGGTGGTGCAGGGCGACCTGGACGTGGGGGGC**<br>**TACGAGGGCGGGCTCGCCGTGAAAGAGTGGCTGCTGGCCCACGAGGGCCACAGACTGGGCAAGCCTGGGCTGGGT**<br>TCCGGACTCAGATCTCGAGCTCAAGCTTCGAATTCTGCAGTCGACGGTACCGCGGGCCCGGGATCCACCGGATCT<br>AGCCACGGGGGTGGCCCCCCCTCGGGGGACAGCGCATGCCCGCTGCGCACCATCAAGAGAGTCCAGTTCGGAGTC<br>CTGAGTCCGGATGAACTGGTAAGCGGCTCTGTCCTCCCCTTCCCCCCTCCTCCCCTGGCGGCGGGGCCGGACGG<br>GGGCTGCGGAAACTTGGCGCTTTCTCGCTGCTTATGGGTGACGGGCCAGGAGCATGGCTCAGCAGCGCCAAGGCC<br>GTGTAGGCGCCAGTCTCGGGCCTCCCAGAGTTATATTTTGCAAAGAGTTGAGCAGGCGTAGCGCTTTGTCCGAGA<br>TGGGGGGCTGGCTGGAGGGTGAGAAGGAGGGAGAGAAGGATGACTGATTTCACACTCCAGAGTTACCGACGTTAA<br>AGGCGATCCGACGAATCCTAGGGAGTCTTTACAAGTTTGGTTAAGAAGCCAAACCCGGGTAGCTTCCTCCTCTTC<br>TGTAATACAAATGTTAACTGAGCACCTGCTCTGTTGAAGACACAGGAGACTCG |

| *TAF1*-Halo HDR Repair template sequence (5’ to 3’) |
| --- |
| ATGGTGTCCGAGCTTGAGAGATGACATGTTTCTCTTCTCAGTTATGGGAGCTATGAGGAGCCTGATCC<br>CAAGTCGAACACCCAAGACACAAGCTTCAGCAGCATCGGTGGGTATGAGGTATCAGAGGAGGAAGAAG<br>ATGAGGAGGAGGAAGAGCAGCGCTCTGGGCCGAGCGTACTAAGCCAGGTCCACCTGTCAGAGGACGAG<br>GAGGACAGTGAGGATTTCCACTCCATTGCTGGGGACAGTGACTTGGACTCTGATGAATCCGGACTCAG<br>ATCTCGAGCTCAAGCTTCGAATTCTGCAGTCGACGGTACCGCGGGCCCGGGATCCACCGGATCTAGCT<br>GGAGCCACCCGCAGTTCGAAAAAACCGGT**GCAGAAATCGGTACTGGCTTTCCATTCGACCCCCATTAT**<br>**GTGGAAGTCCTGGGCGAGCGCATGCACTACGTCGATGTTGGTCCGCGCGATGGCACCCCTGTGCTGTT**<br>**CCTGCACGGTAACCCGACCTCCTCCTACGTGTGGCGCAACATCATCCCGCATGTTGCACCGACCCATC**<br>**GCTGCATTGCTCCAGACCTGATCGGTATGGGCAAATCCGACAAACCAGACCTGGGTTATTTCTTCGAC**<br>**GACCACGTCCGCTTCATGGATGCCTTCATCGAAGCCCTGGGTCTGGAAGAGGTCGTCCTGGTCATTCA**<br>**CGACTGGGGCTCCGCTCTGGGTTTCCACTGGGCCAAGCGCAATCCAGAGCGCGTCAAAGGTATTGCAT**<br>**TTATGGAGTTCATCCGCCCTATCCCGACCTGGGACGAATGGCCAGAATTTGCCCGCGAGACCTTCCAG**<br>**GCCTTCCGCACCACCGACGTCGGCCGCAAGCTGATCATCGATCAGAACGTTTTTATCGAGGGTACGCT**<br>**GCCGATGGGTGTCGTCCGCCCGCTGACTGAAGTCGAGATGGACCATTACCGCGAGCCGTTCCTGAATC**<br>**CTGTTGACCGCGAGCCACTGTGGCGCTTCCCAAACGAGCTGCCAATCGCCGGTGAGCCAGCGAACATC**<br>**GTCGCGCTGGTCGAAGAATACATGGACTGGCTGCACCAGTCCCCTGTCCCGAAGCTGCTGTTCTGGGG**<br>**CACCCCAGGCGTTCTGATCCCACCGGCCGAAGCCGCTCGCCTGGCCAAAAGCCTGCCTAACTGCAAGG**<br>**CTGTGGACATCGGCCCGGGTCTGAATCTGCTGCAAGAAGACAACCCGGACCTGATCGGCAGCGAGATC**<br>**GCGCGCTGGCTGTCGACGCTCGAGATTTCCGGCTGA**GGCTTCCTTTGGGCCTCCTTGGTCAGCCTTCC<br>CTGTTCTCCAGCCTAGGTGGTTCACCTTTCCCCAATTTGTTCATATTTGTACAGTATCTGATCCTGAA<br>ATCATGAAATTAACTAACACCTTAGCCTTTTTAAAAGTAGTAAGTAAATGATAATAAATCACCTCTCC<br>TAATCTTCCTGGGGCAATGTCACCCTTTGATTTAAAACAAAGCAACCCCCTTTCCCCTACCACTACGG<br>AAAAGAGCAAGCTCAT |

**Supplementary Table 2: Homology-directed repair templates for CRISPR-Cas9 knock-in.** Template sequences are annotated as follows: nucleotides highlighted in **green** indicate the upstream homology arm; **purple bold** letters denote the SNAP or HaloTag coding sequence; nucleotides highlighted in **yellow** indicate the downstream homology arm; **green bold** letters mark the start codon; and **red bold** letters indicate the stop codon. The top sequence corresponds to the SNAP-tag knock-in at the *POLR2A* locus, and the bottom sequence corresponds to the HaloTag knock-in at the *TAF1* locus.

| | Sequence (5’ to 3’) |
|---|---|
| SNAP-*POLR2A* forward primer (5’ phosphorylated) | AAGGGGGAGAGACAAACTGC |
| SNAP-*POLR2A* reverse primer | ACAGAAGAGGAGGAAGCTACC |
| *TAF1*-Halo forward primer | GTCCGAGCTTGAGAGATGAC |
| *TAF1*-Halo reverse primer (5’ phosphorylated) | GCTTGCTCTTTTCCGTAGTG |

**Supplementary Table 3: PCR primers to amplify homology-directed repair templates.** The forward primer for amplifying SNAP-*POLR2A* and the reverse primer for *TAF1*-Halo must be 5’-phosphorylated to enable single-strand DNA generation following PCR amplification.

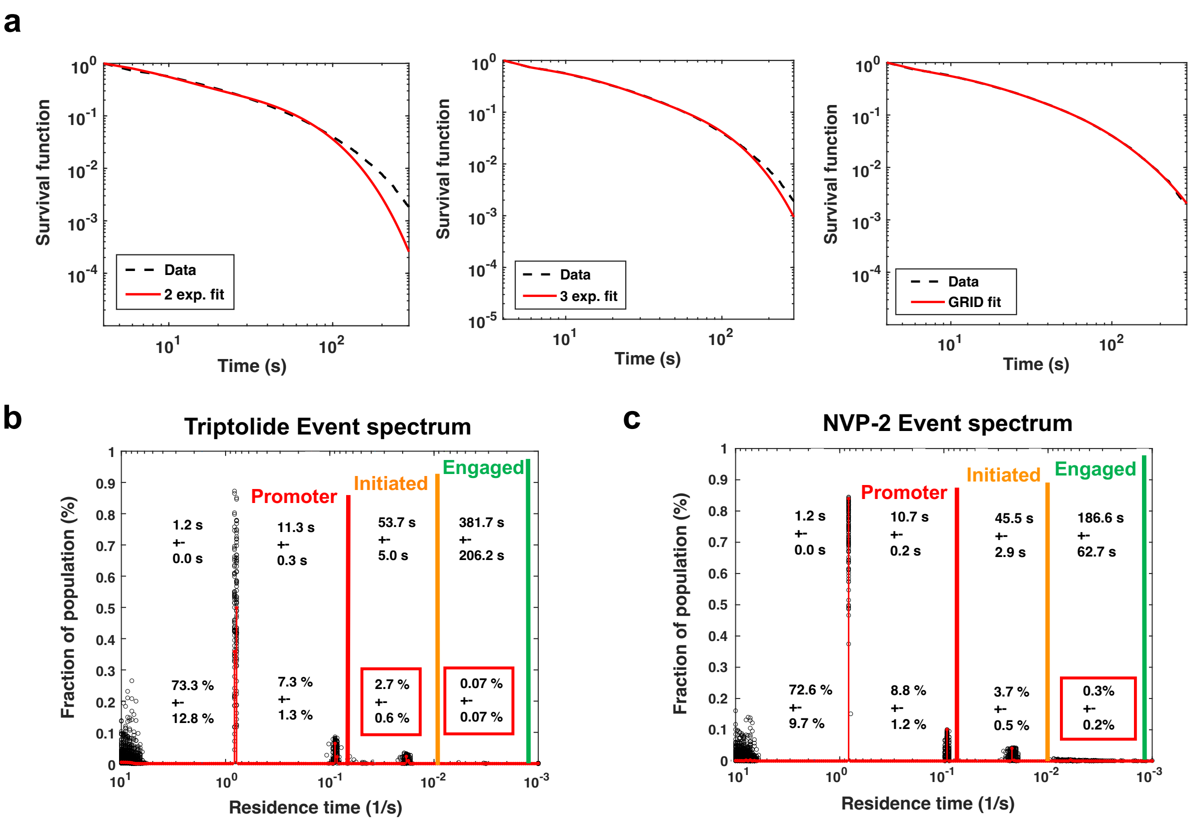


d

| Replicate 1 | $P_1$ (%) | $\tau_1$ (s) | $P_2$ (%) | $\tau_2$ (s) | $P_3$ (%) | $\tau_3$ (s) | $P_4$ (%) | $\tau_4$ (s) |
|---|---|---|---|---|---|---|---|---|
| DMSO (n = 27) | 70.3 ± 11.0 | 1.25 ± 0.02 | 9.6 ± 1.5 | 10.07 ± 0.18 | 4.6 ± 0.8 | 38.2 ± 1.7 | 1.5 ± 0.3 | 121.5 ± 6.1 |
| Triptolide (n = 23) | 73.3 ± 12.8 | 1.25 ± 0.02 | 7.5 ± 1.5 | 11.38 ± 0.35 | 2.7 ± 0.6 | 54.6 ± 2.9 | 0.0 ± 0.0 | 400.0 ± 224.0 |
| NVP-2 (n = 29) | 72.6 ± 9.7 | 1.22 ± 0.00 | 8.76 ± 1.2 | 10.73 ± 0.18 | 3.7 ± 0.5 | 45.5 ± 2.9 | 0.3 ± 0.2 | 186.6 ± 62.7 |

| Replicate 2 | $P_1$ (%) | $\tau_1$ (s) | $P_2$ (%) | $\tau_2$ (s) | $P_3$ (%) | $\tau_3$ (s) | $P_4$ (%) | $\tau_4$ (s) |
|---|---|---|---|---|---|---|---|---|
| DMSO (n = 27) | 65.0 ± 11.3 | 1.27 ± 0.01 | 8.9 ± 1.5 | 10.65 ± 0.20 | 4.0 ± 0.7 | 39.2 ± 2.7 | 1.2 ± 0.3 | 115.7 ± 10.0 |
| Triptolide (n = 25) | 74.5 ± 11.2 | 1.21 ± 0.02 | 6.7 ± 1.0 | 9.70 ± 0.27 | 2.4 ± 0.4 | 43.1 ± 3.8 | 0.3 ± 0.2 | 144.2 ± 54.1 |
| NVP-2 (n = 30) | 64.6 ± 12.2 | 1.22 ± 0.00 | 7.4 ± 1.4 | 10.35 ± 0.20 | 3.1 ± 0.6 | 40.1 ± 2.0 | 0.3 ± 0.1 | 192.4 ± 37.7 |

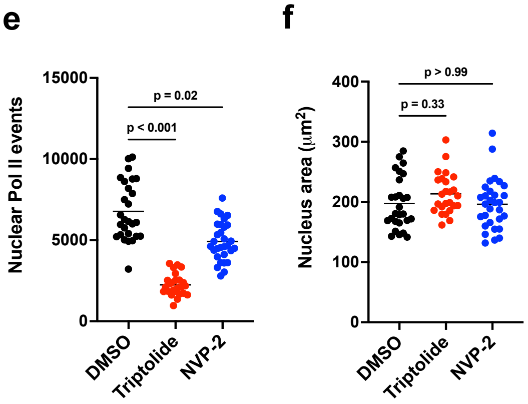


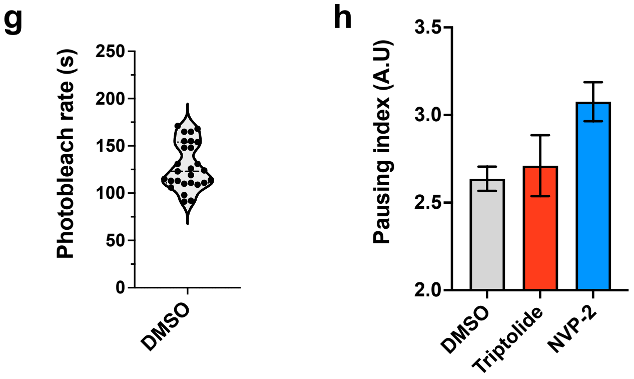

**Supplementary Figure 1: Transcription inhibitors define GRID populations and confirm pausing index trends across replicates. a,** Model fitting comparison of Pol II dwell time distributions using a 2-exponential fit (left), 3-exponential fit (middle), and GRID-fit (right). While the 2- and 3-exponential fits capture some features of the data, the GRID model provides the best overall fit and identifies four distinct kinetic populations. **b,** GRID event spectrum for Pol II binding in cells treated with 1 µM Triptolide. Red boxes indicate changes in populations relative to control (DMSO) across two separate days of imaging. **c,** GRID event spectrum for cells treated with 50 µM NVP-2. Red boxes indicate changes in populations relative to control (DMSO) across two separate days of imaging. **d,** Summary tables of the mean population fraction and residence time of the four kinetic Pol II populations across DMSO, Triptolide, and NVP-2 treatment conditions. Top table (pink) corresponds to replicate 1 (Day 1); bottom table (blue) shows replicate 2 (Day 2). Cells shaded in red indicate a decrease in population fraction or residence time relative to DMSO for that replicate; green indicates an increase. **e,** Total number of nuclear Pol II binding events per nucleus in DMSO (n = 27), Triptolide (n = 25), and NVP-2 (n = 30) treatment conditions from replicate 2. **f,** Total nuclear area per nucleus in DMSO (n = 27), Triptolide (n = 25), and NVP-2 (n = 30) treatment conditions from replicate 2. **g,** Photobleach rates per cell in DMSO-treated nuclei from replicate day 1. Rates were estimated by measuring the total nuclear fluorescence decay over time and fitting each trace to a single-exponential decay model. The average photobleach rate across cells was ~128 seconds. **h,** Pausing index values for replicate 2 (Day 2), calculated as the number of molecules with residence times between 40–90 seconds (paused) divided by those >90 seconds (elongating). Poisson-based error estimates are shown as mean ± SEM.

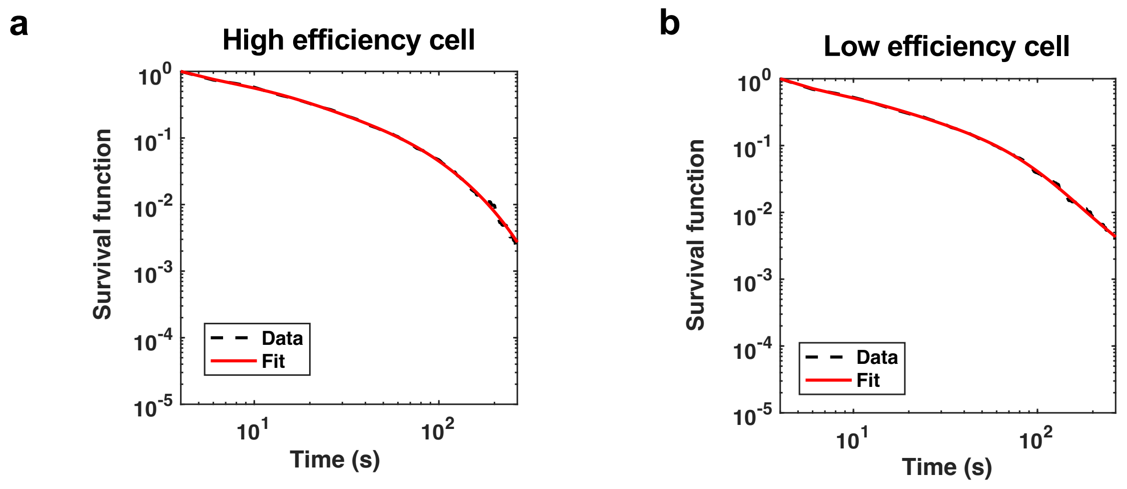


**Supplementary Figure 2: Supporting data for single-cell analysis of Pol II transcription efficiency**. **a,** GRID-based model fitting of Pol II dwell time distribution for the high-efficiency DMSO-treated nucleus shown in Fig. 3a. **b,** GRID-based model fitting of Pol II dwell time distribution for the low-efficiency DMSO-treated nucleus shown in Fig. 3b.

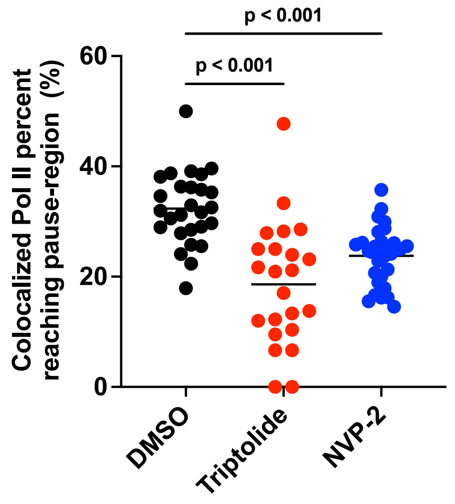


**Supplementary Figure 3: Drug treatment decreases the fraction of colocalized Pol II reaching the pausing region**. Initiation efficiency of colocalized Pol II molecules under DMSO, Triptolide, and NVP-2 treatment, defined as the percentage of molecules with residence times ≥ 40 seconds. Statistical significance assessed using a Kruskal-Wallis test.
